# Structural basis of bile acid receptor activation and Gs coupling

**DOI:** 10.1101/2020.05.24.104034

**Authors:** Fan Yang, Chunyou Mao, Lulu Guo, Jingyu Lin, Qianqian Ming, Peng Xiao, Xiang Wu, Qingya Shen, Shimeng Guo, Dan-Dan Shen, Ruirui Lu, Linqi Zhang, Shenming Huang, Yuqi Ping, Chenlu Zhang, Cheng Ma, Kai Zhang, Xiaoying Liang, Yuemao Shen, Fajun Nan, Fan Yi, Vincent C Luca, Jiuyao Zhou, Changtao Jiang, Jin-Peng Sun, Xin Xie, Xiao Yu, Yan Zhang

## Abstract

G protein-coupled bile acid receptor (GPBAR) is a membrane receptor that senses bile acids to regulate diverse functions through Gs activation. Here, we report the cryo-EM structures of GPBAR–Gs complexes stabilized by either high-affinity P395 or the semisynthesized bile acid derivative INT-777 at 3-Å resolution. These structures revealed a large oval-shaped ligand pocket with several sporadic polar groups to accommodate the amphipathic cholic core of bile acids. A fingerprint of key residues recognizing diverse bile acids in the orthosteric site, a putative second bile acid binding site with allosteric properties and structural features contributing to bias property were identified through structural analysis and mutagenesis studies. Moreover, structural comparison of GPBAR with other GPCRs uncovered an atypical mode of receptor activation and G-protein– coupling, featuring a different set of key residues connecting the ligand binding pocket to the Gs coupling site, and a specific interaction motif localized in intracellular loop 3. Overall, our study not only provides unique structural features of GPBAR in bile acid recognition, allosteric effects and biased signaling, but also suggests that distinct allosteric connecting mechanisms between the ligand binding pocket and the G protein binding site exist in the GPCR superfamily.

## Introduction

Bile acids are important endocrine and amphipathic signaling molecules that are synthesized from cholesterol in the liver and further diversified by the gut microbiota^1,2^. Their diverse biological effects on mediating insulin resistance, obesity, lipid metabolism, and systemic metabolic control are exerted in conjunction with the nuclear farnesoid X receptor (FXR) and the membrane-bound G protein-coupled bile acid receptor GPBAR (TGR5 or GPR131)^3^. GPBAR has been found in a wide range of tissues and serves as a signaling hub in the liver–bile-acid–microbiota–metabolism axis^1,3,4^.

Bile acids induce both beneficial and adverse effects in different pathophysiological conditions via GPBAR. For example, cholic acid (CA) and taurocholic acid (TCA) increase energy expenditure and reduce adiposity through activation of GPBAR^5,6^. Tauroursodeoxycholic acid (TUDCA) has been used in traditional Chinese medicine for more than 3000 years and shows anti-inflammatory effects in the liver and promotes nitric oxide (NO) release and vasodilation in the heart^1,7^. Conversely, lithocholic acid (LCA) has been reported to cause insulin resistance, and deoxycholic acid (DCA) has been shown to promote cancer cell progression. Apart from the differences in distribution and metabolism among bile acids, the diverse downstream pathways of GPBAR also contribute to various functional outcomes. Many of the beneficial effects of bile acids, such as protecting against obesity and diabetes, combating steatosis and reducing inflammation^8-12^, have been attributed to GPBAR–Gs coupling. In addition, GPBAR signals to β-arrestin to activate SRC kinase and induce innate antiviral immune response in divergent cell types^13,14^, Notably, sequence alignment of GPBAR with other family A GPCRs whose structures are available implies that it lacks the conserved NPxxY motif and has a shortening at intracellular loop 2 (ICL2), indicating a potential different activation mechanism (Extended Data Fig. 1)^15,16^. Due to the paucity of knowledge about the detection of amphipathic ligands by membrane receptors, the diversity in function and signalling after the engagement of GPBAR with different bile acids, the lack of several conserved motifs required for GPCR activations, there is an urgent need to delineate the molecular mechanism underlying GPBAR activation in response to various bile acids. Here, we determined 3-Å cryo-EM structures of the GPBAR–Gs in complexes with P395 and INT-777, a highly potent synthetic agonist and a semisynthesized bile acid derivative with beneficial effects in nonalcoholic steatohepatitis (NASH) in preclinical animal studies, respectively. These structures, provide key knowledge for an unconventional activation mechanism of GPBAR in response to agonists, a detailed fingerprint for the recognition of diverse bile acids, the structural basis for biased GPBAR signalling, an alternative GPCR**–**Gs**–**protein engagement mode and a potential second bile acid binding site with allosteric properties.

**Figure 1.**
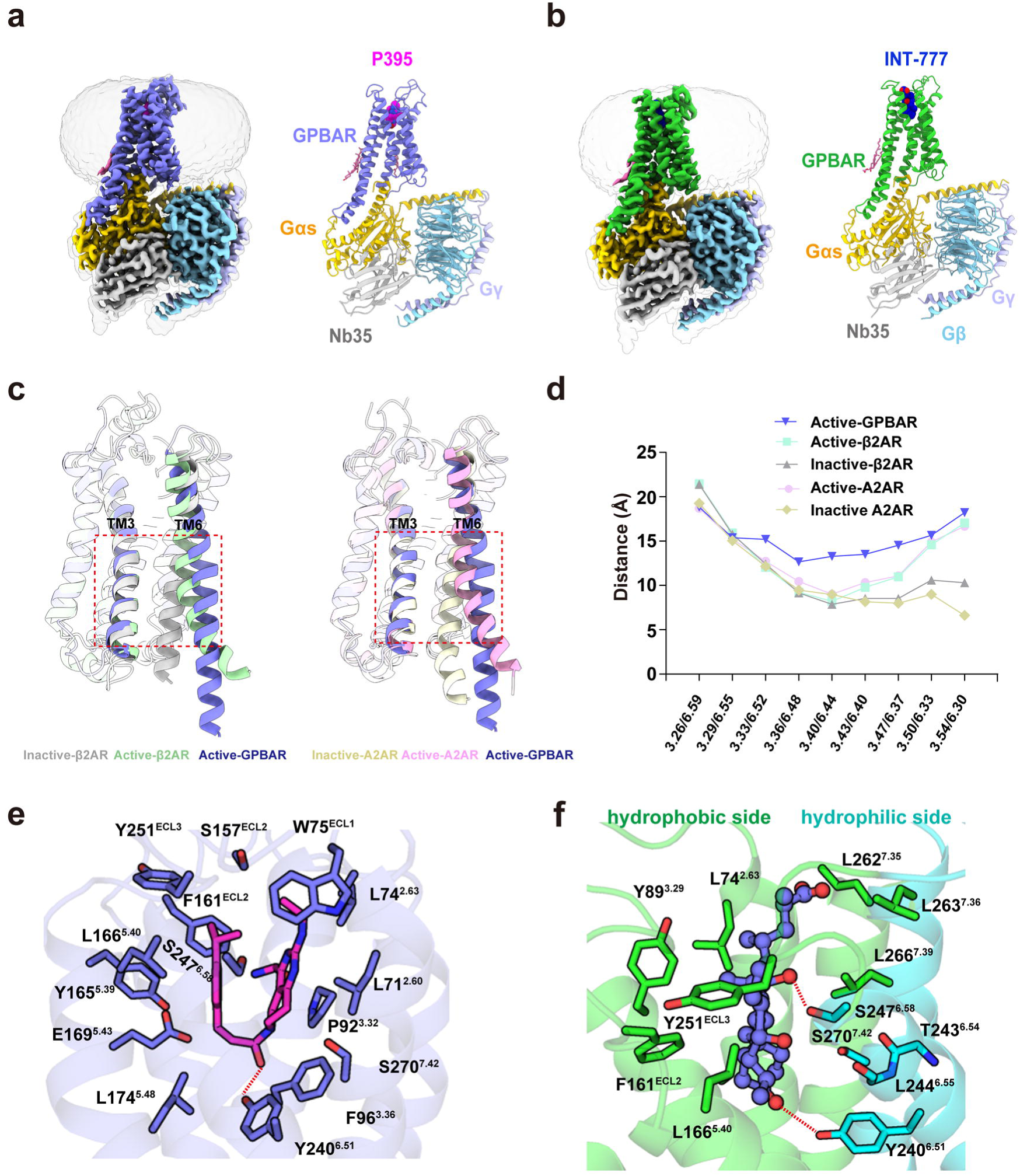
Cryo-EM structure of 395–GPBAR–Gs and INT-777–GPBAR–Gs complex. **a-b** Cryo-EM density (left panel) and ribbon representation (right panel) of the P395**–** GPBAR**–**Gs complex (a) or INT-777–GPBAR–Gs complex (b). P395, magentas; GPBAR (a), slate; Gαs, yellow; Gβ, cyan; Gγ, light blue; Nb35, gray; INT-777, blue; GPBAR (b), green. **c**, Ribbon representation of the larger separation of TM3 and TM6 in active GPBAR compared to that in active β2AR (PDB ID 3SN6) in complex with Gs, inactive β2AR stabilized by an antagonist (PDB ID 3NYA), active A2AR in complex with miniGs (PDB ID 5G53) and inactive A2AR (PDB ID 3EML). **d**, Plot of Cα distances of residues between TM3 and TM6 of active GPBAR, active/inactive A2AR and active/inactive β2AR. **e**, Structural view of the insertion of P395 into the ligand pocket composed of residues from TM2, TM3, TM6 and TM7 and enclosed by three extracellular loops. The hydrogen bond is depicted as a dashed line. A notable feature of the interactions between P395 and GPBAR is their hydrophobic nature, with ten hydrophobic residues involved and only one polar contact. **f**, Detailed interactions between INT-777 and the GPBAR. Hydrogen bonds are highlighted with red dashes.

## Results

### Complex formation and cryo-EM analysis

Full-length human GPBAR with thermostabilized cytochrome b_562_RIL (BRIL) introduced into the N-terminus was co-expressed with Gs protein in *Spodoptera frugiperda* (Sf9) insect cells. Active complexes were readily formed by the addition of excess high-affinity agonist P395^17^ and the nanobody Nb35 ^16^ (Extended Data Fig. 2a), however, the low solubility and affinity of endogenous bile acids complicate the formation of bile acid–GPBAR–Gs complexes *in vitro*. We screened a panel of bile acids and identified that only INT-777 robustly promoted a high fraction of GPBAR–Gs complex formation (Extended Data Fig.2b,). The GPBAR–Gs complexes stabilized by P395 or INT-777 were purified and analysed by single-particle Cryo-EM, which enabled us to construct electron density maps with an overall resolution of 3.0-Å (Extended Data Fig. 2c-2f, Extended table 1). Atomic resolution structures of GPBAR including all seven transmembrane (TM) helices with both intracellular and extracellular loops (ICLs and ECLs, respectively) were confidently modelled using the high-resolution electron density, and the majority of the side chains from the receptor and the G proteins were clearly identified (Fig. 1a-1b and Extended Data Fig. 3). In particular, a well-defined density was observed for ICL3, which is not well resolved in any of the available GPCR**–**Gs structures (Fig. 1a-1b and Extended Data Fig. 3c, 3d). In the receptor orthosteric binding pocket, well-defined electron densities were unambiguously assigned to the compound P395 or the bile acid derivative INT-777 (Extended Data Fig. 3c, 3d)

**Figure 2.**
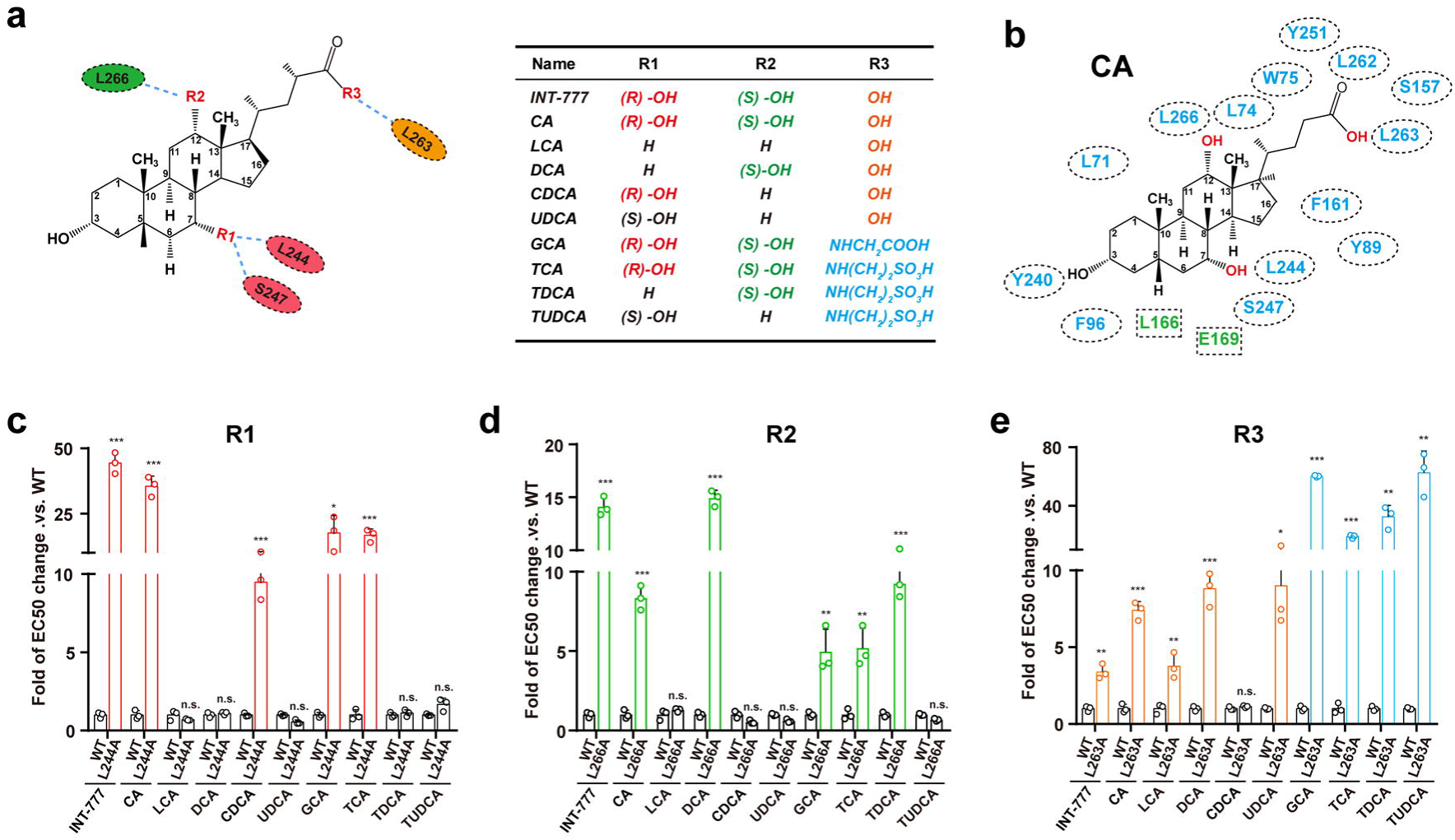
Structural fingerprints of GPBAR recognizing different bile acids. **a**, Diagram of the fingerprint that differentiates diverse bile acids (left panel). The 7 (R1), 12 (R2) and C-terminal (R3) positions are the most common substitution or conjugating sites in the primary bile acid CA to generate diverse bile acids, which are shown in red. Residues shown for interaction with the R1, R2 and R3 positions in GPBAR are shaded in red, green, and yellow, respectively. Substitution and conjugation status of INT-777, CA and several different bile acids at the R1, R2 and R3 positions are summarized in a table shown on the right panel. **b**, Diagram of the potential primary bile acid Cholic Acid (CA) interaction in the ligand binding pocket of GPBAR. Blue, residues located in the INT-777 binding pocket and shown mutating effects on both CA and INT-777; Green, residues with mutating effects only on INT-777, but not CA. The mutating effects were referred to Extended data 5c-d. **c**-**e**, Effects of bile acid recognition fingerprint mutants on cAMP accumulation induced by different bile acids. (**c**), mutation of L244 to A; (**d**), mutation of L266 to A; (**e**), mutation of L263 to A. The fold of EC50 change of mutant.vs. wild type for each individual bile acid were used for straightforward view. The original data were referred to Extended data 6. Values are the mean ± SEM of three independent experiments for the wild type (WT) and mutants. Statistical differences between WT and mutations were determined by One-way ANOVA (**, P<0.01; ***, P<0.001, n.s., no significant difference)

**Figure 3.**
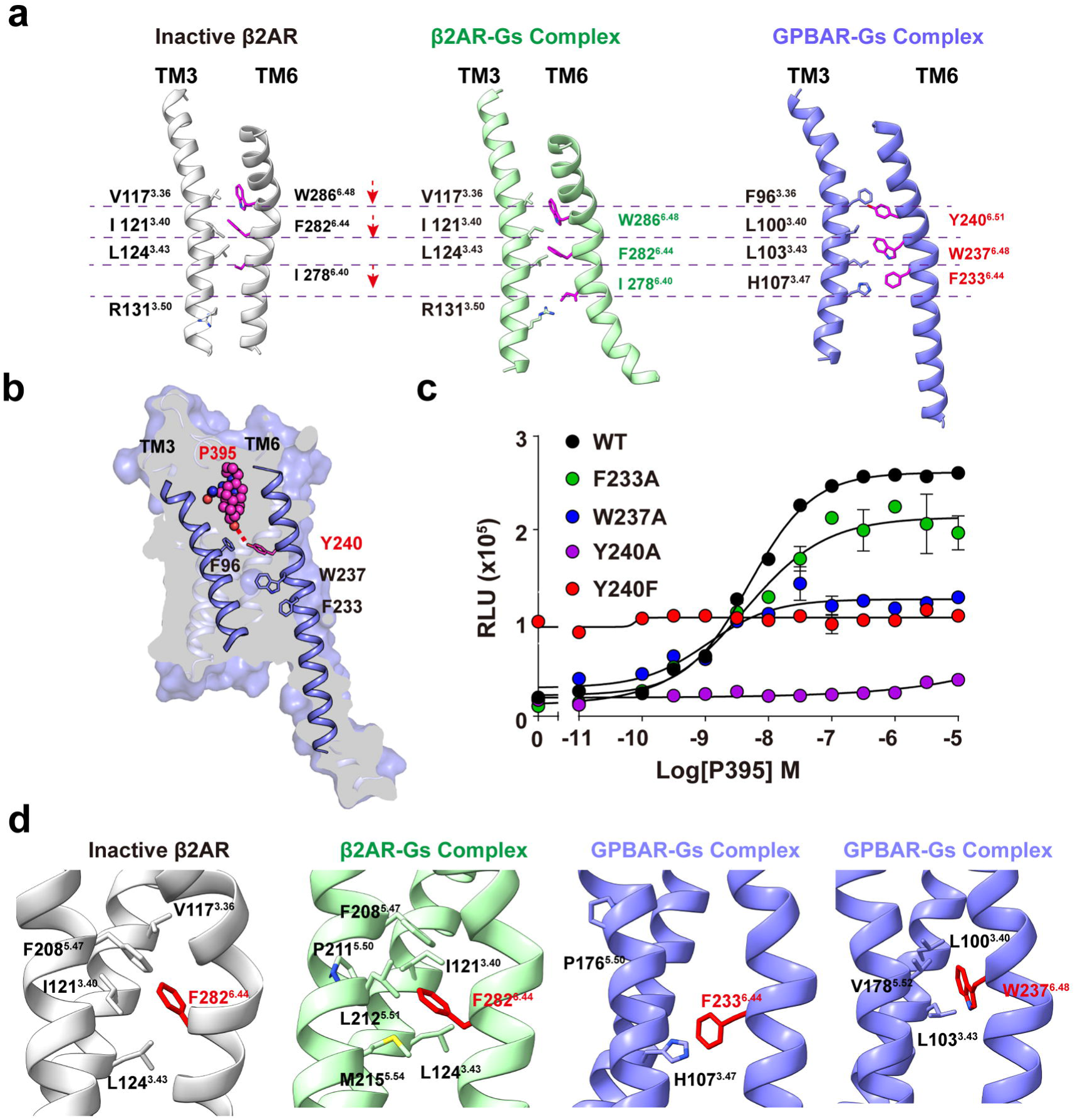
An unexpected activation mechanism of GPBAR. **a**, Structural representation of the important residues participating in GPBAR activation, including Y240^6.51^, the presumed toggle switch W237^6.48^, the L100^3.40^ and F233^6.44^ of the P^5.50^I^3.40^F^6.44^ motif, and compared them with their counterparts in the inactive β2AR (PDB ID 3NYA) and the active β2AR in complex with the agonist BI and Gs (PDB ID 3SN6). Notably, Y240^6.51^ of GPBAR assumes the same position as W286^6.48^ in the β2AR-Gs complex, which undergoes a downshift of one helical turn in relation to TM3 during the transition from the inactive to the active state. **b**, Cutaway view of key residues governing GPBAR activation in response to P395 binding. **c**, Dose response curves of GPBAR carrying mutations in the key residues involved in activation in the cAMP accumulation assay in response to P395. Whereas the F233A has very little effect on P395 induced cAMP accumulation, the Y240A and Y240F totally eliminated the response to P395 engagement. There is a significant high level basal activity of the Y240F mutant. Data are shown as mean ± SEM from three independent measurements. **d**, Lack of the compact structural P^5.50^I^3.40^F^6.44^ motif in GPBAR structure. Left, structural rearrangement of the PIF motif during β2AR activation. Right, separation of P^5.50^L^3.40^F^6.44^ in the GPBAR structure. Instead, W237^6.48^ forms hydrophobic interactions with L100^3.40^ and V178^5.52^ to constitute a VLW motif in GPBAR.

Although significant differences were observed in the ligand binding mode, extracellular motifs and a putative second bile acid binding pocket, the overall architecture of the INT-777– GPBAR–Gs complex is very similar to that of the P395–GPBAR–Gs complex. Three distinct yet intercorrelated features were observed for GPBAR–Gs complexes when comparing active GPBAR with those of other class A receptor**–**Gs complexes, including (1) TM6 had an overall larger separation from the central TM3 (Fig. 1c-1d), (2) the C-terminal end of the α5 helix did not penetrate so deeply into the receptor 7-TM core as other Gs-coupled structures, and (3) ICL3 was specifically coupled to the Gs protein.

### Binding of P395 and INT-777 in the orthosteric site

GPBAR expands a large ovate pocket to accommodate the bulky P395 ligand or bile acids. Inside the orthosteric pocket, a long-stranded hydrophobic strip from TM2 and TM3 holds the tetrahydropyrido[4,3-d] pyrimidine moiety, whereas another hydrophobic patch from TM5 accommodates the 4-isopropylphenyl part of P395 (Fig. 1e and Extended Data Fig. 4a,4b). The restraint of these sidewalls forces the folding of P395 into a U-shaped configuration rather than an extended topology, as predicted by previous studies ^18^ (Fig. 1e and Extended Data Fig. 4b,). In contrast to P395, INT-777 assumes a shovel-like structural conformation with folding between ring A and the rest of the steroid core (Fig. 1f and Extended Data Fig. 4c, 4d). The cyclopentano-bicyclohexyl part (ring B-C-D) is inserted into the ligand binding pocket of GPBAR in a direction parallel to TM2 (Fig. 1f and Extended Data Fig. 4d, 4e). Interestingly, there is a 90-degree difference between the face of the INT-777 steroid nucleus core and the face of P395, indicating that ligands with distinct chemical features could be accommodated by the ligand binding pocket of GPBAR (Extended Data Fig. 4f).

**Figure 4.**
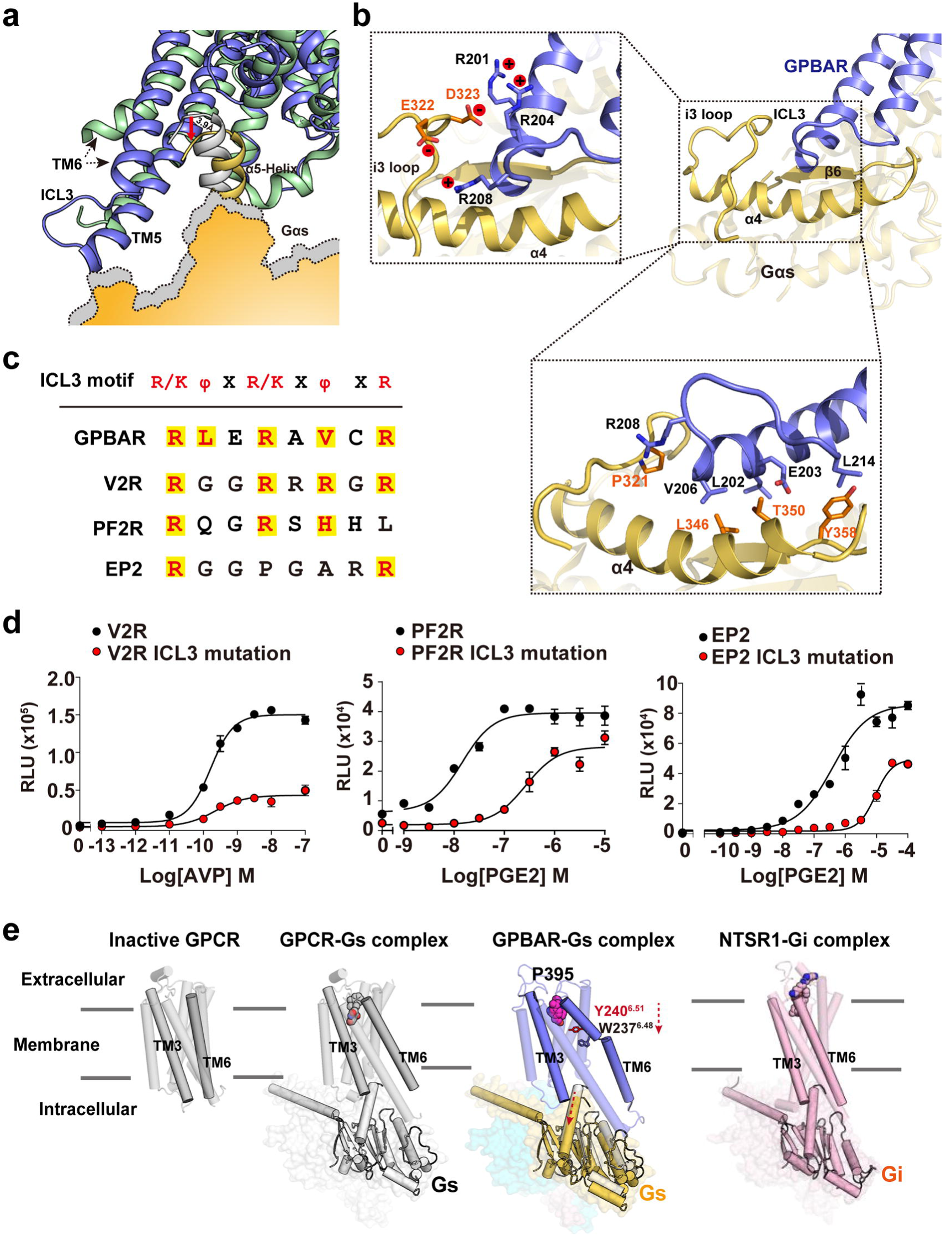
The coupling of GPBAR to Gs. **a**, Schematic representation of the downshift of the α5 helix of the Ras-like domain of Gs, which is potentially due to the longer extension of TM6, the rigidity of ICL3 of GPBAR and its strong interaction with Gs. Ribbon representation: GPBAR, slate; β2AR, green; α5-helix of Gs bound to β2AR, grey; α5-helix of Gs bound to GPBAR, yellow. Surface representation: Gs bound to β2AR, grey; Gs bound to GPBAR, yellow. **b**, Specific interactions of the ICL3 of GPBAR with the Ras-like domain of Gs. An overall view of GPBAR ICL3 and Gs interaction are shown on right upper panel. The ICL3 of GPBAR, i3-loop, β6 and α4 of Gs are highlighted. Specific charge interactions and hydrophobic interactions (lower right panel) are depicted between the interface of the GPBAR ICL3 and Gs. **c**, Sequence comparisons of several known Gs–coupled GPCRs that have similar residues to the R/KΨXR/KXΨXR motif in ICL3, including GPBAR, V2R, PF2R and EP2. Residues that are in consistent with this motif are shaded with yellow. **d**, Effects of ICL3 mutations in the R/KΨXR/KXΨXR motifs of V2R (R243A/R247A/R249A/R251A), PF2R (R238A/R241A/HR243A) and EP2 (R242A/R249A) on their agonist-induced cAMP accumulation. Data are shown as mean ± SEM from three independent measurements. **e**, A cartoon model illustrating the structural differences of the activation and Gs coupling of GPBAR compared to the other class A GPCR**–**Gs or GPCR**–**Gi complexes. From the left to right is the inactive GPCR structural model (using β2AR as an example, PDB ID 3NYA), the general GPCR**–**Gs complex (using β2AR as an example, PDB ID 3SN6), the GPBAR**–**Gs complex and the NTSR**–**Gi complex (PDB ID 6OS9). Compared to other class A GPCR**–**Gs complexes or NTSR**–**Gi complex, the GPBAR**–**Gs complex exhibits distinct features, first a larger separation at the TM3-TM6 helices in the center of receptor region, second the H5 of Gs in GPBAR**–**Gs complex showing one helical turn downshifting probably due to the direct interaction of the ICL3 of GPBAR with the Gs.

Seven hydrophobic residues and three polar residues inside the GPBAR orthosteric site form common interactions with both P395 and INT-777 (Fig. 1e-1f, Extended Data Fig. 5a and Extended Table 2, 3). Mutation of these hydrophobic residues to alanine significantly impaired both P395 and INT-777 interactions (Extended Data Fig. 5b, 5c). Unlike the hydrophobic nature of P395, one unique characteristic of INT-777 and other bile acids is amphipathic, with all hydroxyl substituents and a C-terminal carboxyl group pointing to one side, leaving a convex hydrophobic surface on the other side (Fig. 1f). The convex surface of the INT-777 hydrophobic side faces toward TM5, ECL2 and ECL3, forming extensive interactions between the rings B, C and D and the aromatic residues. On the hydrophilic side of INT-777, the hydroxyl groups of Y240^6.51^, S247^6.58^, and S270^7.43^, as well as the backbone carboxyl of T243^6.54^, L244^6.55^ and S247^6.58^, provide a sporadic polar environment (superscripts referring to Ballesteros-Weinstein number^19^, Fig. 1f).

**Figure 5.**
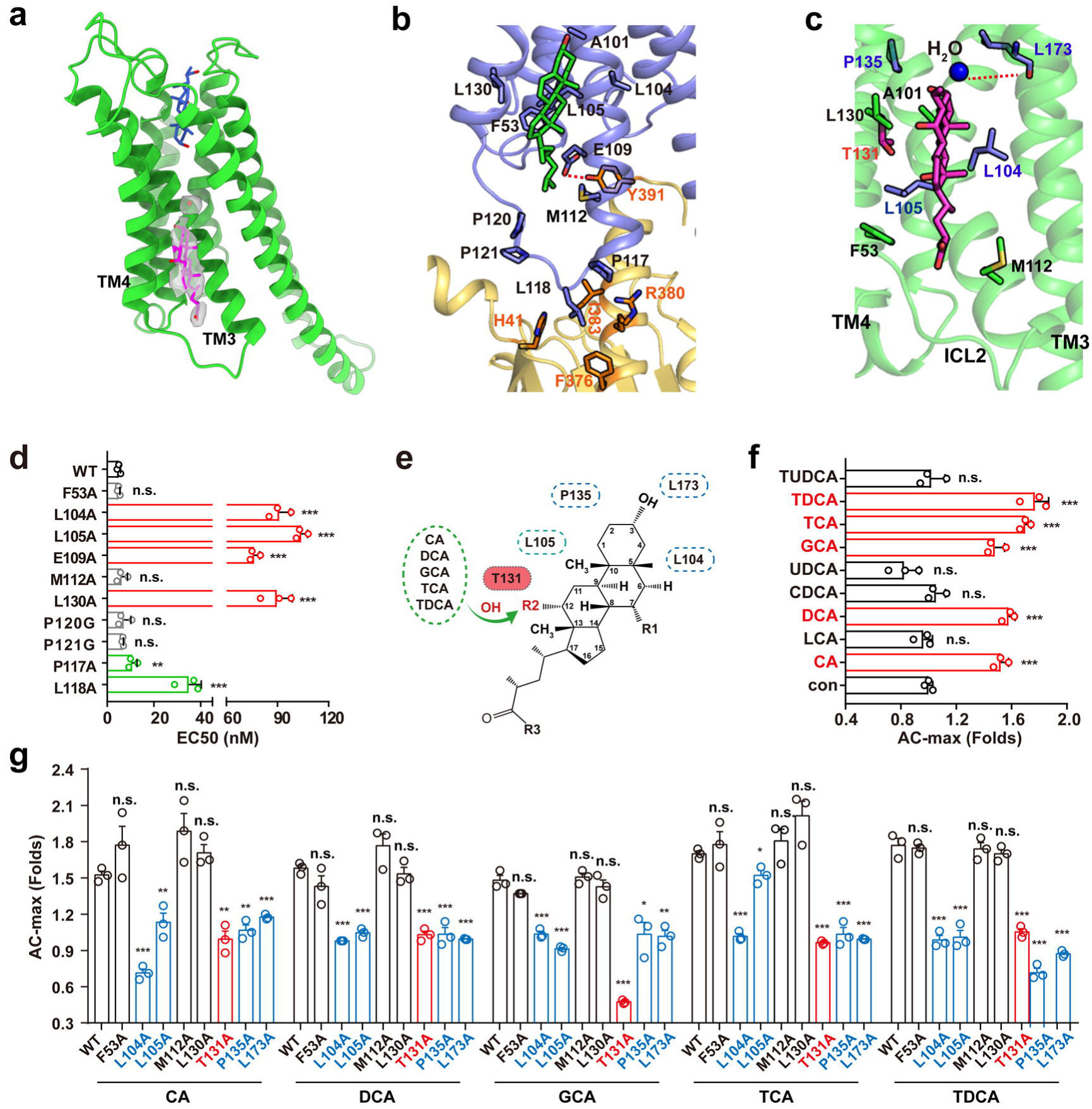
The second ligand binding pocket and its allosteric effect. **a**, A cartoon presentation of GPBAR complex highlighting the existence of a potential second ligand binding pocket of GPBAR. Upper, an INT-777 bound to the orthosteric site, Lower left, an INT-777 molecular binds to an allosteric site. The two sites are mainly connected by TM3. **b**, Possible interactions between the modelled cholesterol with TM2, TM3, TM4 and TM5 of the receptor. Residues constituted the second ligand binding site (side chains located within 4Å between the modelled P395 and the GPBAR) are highlighted in stick. This model was used for further mutagenesis validation. **c**, Possible interactions between the modelled second INT-777 with TM2, TM3, TM4 and TM5 of the receptor. Residues constituted the second ligand binding site (side chains located within 4Å between the modelled INT-777 and the GPBAR) are highlighted in stick. The model was then used for mutagenesis evaluation. **d**, Effects of different second binding pocket mutations on the efficacy of P395-induced cAMP accumulation. Values are the mean ± SEM of three independent experiments for the wild type (WT) and mutants. Statistical differences between WT and mutations were determined by One-way ANOVA (**, P<0.01; ***, P<0.001, n.s., no significant difference) **e**, Diagram of the potential interacting mode of bile acid within the allosteric ligand binding pocket of GPBAR. Five bile acids, including CA, DCA, GDA, TCA and TDCA who share the common 12-OH substitution, engaged with T131 in the second ligand binding pocket, which is essential for the allosteric effects. Other upper four residues, including L104, L105, P135 and L173 in the second ligand binding pocket, also mediates the allosteric effects. **f**, Allosteric effects of different bile acids toward P399 induced cAMP accumulation. The max of allosteric cooperativity (AC-max) derived from the dose response curve was shown. The original data is referred to Extended data table 9. **g**, The effects of mutations of residues in second ligand binding pocket on the allosteric effects of different bile acids. The original data was referred to Extended table 10. d, f, g: EC50 values or Allosteric cooperativity max are the mean ± SEM of at least 3 independent experiments. Statistical differences between WT and mutations were determined by One-way ANOVA (*, P<0.05; **, P<0.01; ***, P<0.001, n.s., no significant difference).

At the bottom of the orthosteric site, both the shovel structure of INT-777 and the U-shaped configuration of P395 facilitated the seating of the shovel head or the acyl linker, respectively, into a hydrophobic pocket cleft, with the strong hydrophobic packing interaction with F96^3.36^ (Extended Data Fig. 5e, 5f). The 3-OH substituent of the steroid nucleus core of INT-777 and the oxygen of the acyl linker of P395 engage in hydrogen bonding (H-bonding) with the hydroxyl group of Y240^6.51^ (Fig. 1e-1f and Extended Data Fig. 5e, 5f). Both the Y240F and S247A mutants displayed significantly increased constitutive activity but only very weak induced activation in response to either INT-777 or P395 engagement (Extended Data Fig. 5g, 5h), indicating that the intramolecular polar network involving S247^6.58^ and Y240^6.51^ may be required to maintain GPBAR in an inactive state.

### Structural fingerprints of GPBAR recognizing bile acid

Endogenous bile acids have the same core but are differentiated mainly by hydroxylation at the 7 (R1) and 12 (R2) positions of the 23-carbon steroid nucleus or by amidation at the carboxyl terminus (R3) (Fig. 2a). All tested bile acids activate GPBAR but with different potencies and efficacies for inducing cAMP accumulation^1^, indicating that these compounds have different abilities for inducing Gs coupling with GPBAR, most likely due to the distinct receptor conformations stabilized by the corresponding bile acids. These GPBAR conformational differences may not only affect Gs signalling but also contribute to the functional diversity of bile acids through arrestin or other downstream effector proteins. Therefore, generalizing the principle underlying the interaction between various bile acids and GPBAR is crucial for the selective usage of bile acid derivatives to treat human diseases.

The structure of the INT-777–GPBAR complex provided a starting model for investigating the interaction mode of other bile acids within GPBAR. Ligand binding and mutagenesis scanning identified 13 residues of GPBAR that are common interaction residues for both INT-777 and CA, a primary native bile acid (Fig. 2b and Extended Figure 5d). Mutations of L166 and E169 only affected INT-777, likely due to the ethyl group at the 6 position of INT-777 (Fig. 2b and Extended Figure 5d). These results suggest that CA shares a very similar binding mode of INT-777 and that the INT-777-GPBAR complex is a useful model for studying interactions between GPBAR and bile acids. We next investigated the specific residues responsible for recognizing the hydroxyl groups attached to the R1 or R2 positions and the conjugating groups at its carboxyl terminus, which are mostly diversified in different bile acid structures and could be determinants of their various biological activities (Fig. 2a). The R1 position (R)-OH forms a hydrogen bond with S247^6.58^ and a hydrophobic interaction with L244^6.55^. The hydroxyl group at the R2 position participates in hydrophobic interactions with L266^7.39^. Finally, the carboxyl tail of INT-777 forms specific contact with L263 (Fig. 2a).

Due to the weak binding of several bile acids, which poses a great challenge for binding assays, we used the cAMP assay to functionally examine the mutagenesis effects of these potential key bile acid interacting residues toward all 9 commercially available bile acids (Fig. 2c-2e; Extended Data Fig 6). Specifically, the L244A mutation significantly decreased the half-maximal effective concentration (EC50) of INT-777, CA, CDCA, GCA and TCA, which all have a hydroxyl group at the R1 position. In contrast, the effect of L244A on other tested bile acids, including LCA, DCA, UDCA, TDCA and TUDCA, showed no significant effect (Fig. 2c). The mutating effects of L266A also paired well with the bile acids that had a hydroxyl group at the R2 position (Fig. 2d). Intriguingly, the L263A mutation moderately decreased the Gs activity of GPBAR in response to the engagement of bile acids that had a hydroxyl group at the R3 position, such as LCA, DCA and UDCA, but it had much larger effects on the EC50 of cAMP accumulation elicited by GCA, TCA, TDCA and TUDCA, all of which had larger groups conjugated at the R3 position at the end of the steroid core (Fig. 2e, Extended Data Fig. 6). In summary, the combination of the mutating effects and the INT-777–GPBAR complex structure revealed that triplet leucine cluster (L244^6.55^, L263^7.36^ and L266^7.39^), as well as a potential role of S247^6.58^, constitute a fingerprint reader to discriminate the interactions between different bile acids and GPBAR (Fig. 2a).

**Figure 6.**
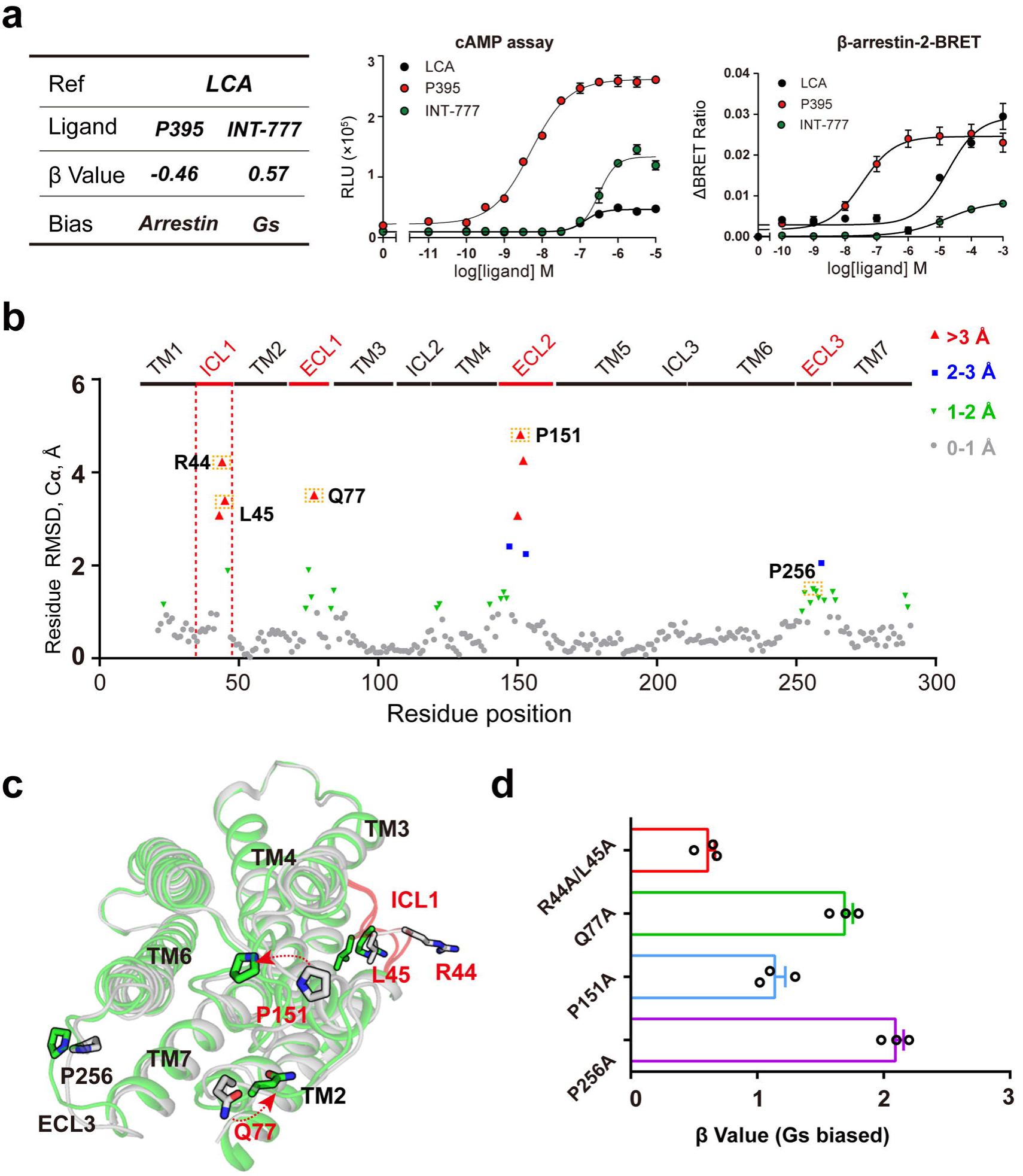
Structural basis of the biased agonism by INT-777. **a**, Comparison of the biased properties of INT-777 and P395. Both INT-777 and P395 were assessed for cAMP signalling (left panel) and β-arrestin-2 recruitment (middle panel). The bias factor (β value) of P395 was calculated using a native bile acid LCA as the reference. The p395 is a β-arrestin-2 biased ligand with respect to INT-777. β > 0 indicates Gs biased, β < 0 indicates arrestin biased. The significant negative β value clearly indicates that P395 is a β-arrestin-biased ligand. Data from three independent experiments are presented as mean ± SD. **b**, Plot of the distance root-mean-square deviations (RMSDs) of each residue between INT-777–GPBAR and P395–GPBAR structures. The horizontal and vertical axes indicate the amino acid sequence of the GPBAR and the RMSDs (Cα deviations) for every residue, respectively. The red, blue, green and grey dots represent Cα deviations that range from >3, 2∼3, 1∼2 or <1, respectively. **c**, Extracellular view of the GPBAR transmembrane bundle showing the location of the residues with different RMSD between INT-777–and P395–bound GPBAR, coloured in green and grey respectively. Residues with significant conformational changes, including Q77 and P151, as well as the potential arrestin interaction sites R44 and L45 are highlighted in red. **d**, Biased property analysis of the residues highlighted in (**c**). β values calculated from the molecular efficacies of P395. Positive β values denote Gs-biased signalling using WT GPBAR as the reference. β values are calculated from at least 3 independent experiments.

### An unconventional activation mechanism

A unique characteristic of the activated GPBAR is located in the TM5-ICL3-TM6 region, featuring the more contracted intercellular rim of TM6 and the overall loose contact between TM3 and TM6 in the middle of the TM region compared with those of other activated class A GPCRs (Fig. 1b, 1c and 3a) ^15,20^_,21_. The intracellular end of TM6 of GPBAR is displaced outwards from the receptor core to a similar extent to that in GPCR**–**G_i_ or G_11_ structures rather than GPCR**–**G_s_ complex structures (Extended data Fig. 7a-7c)^15,22-26^. The difference in the TM6s between GPBAR and other active receptors begins at Y240^6.51^, a critical residue that recognizes the core scaffold of both P395 and the semisynthetic bile acid derivative INT-777, and propagates the binding signal through helix turns that enclose the residues from W237^6.48^ to Q222^6.33^ (Fig. 1d, Fig. 3a, 3b). Sequence alignment shows that GPBAR contains the conserved toggle switch W237^6.48^ and proline kink P176^5.50^ (Extended Data Fig. 7d); however, these features do not assume the same positions as presented in the β2AR**–**Gs or A2A**–**Gs complex structures (Fig. 3a, Extended Data Fig. 7e). In both GPBAR**–**Gs complex structures, W237^6.48^ of GPBAR is one helical turn lower than W286^6.48^ of β2AR or W246^6.48^ of A2A in their active conformations (Fig. 3a, Extended Data Fig.7f). In the active structures of the β2AR**–** Gs complex or the A2AR**–**Gs complex, when compared with their inactive states, the hydrophobic interactions between the agonist and the toggle switch W^6.48^ forced TM6 to move one step downward relative to TM3. This shift enabled W286^6.48^ of the active β2AR to form new hydrophobic interactions with V117^3.36^ and I121^3.40^. However, W286^6.48^ of β2AR was substituted with Y240^6.51^ of GPBAR in the same position (Fig. 3a, Extended Data Fig. 7f). GPBAR Y240^6.51^ donates a hydrogen bond to the bound agonists, and undergoes hydrophobic stacking with F96^3.36^ (Fig. 3b), which recalls the functions of the ‘twin-toggle-switch’ of the CB1 receptor^22,27^. Consistently, mutation of Y240F shows no response to P395 engagement and Y240A completely eliminated P395, INT-777 or other bile acid-induced cAMP accumulation (Fig. 3c and Extended data 7g). Collectively, the combination of structural and biochemical analyses suggests that Y240^6.51^ is the functional “toggle switch” of the GPBAR, rather than the conventional W237^6.48^ predicted from the sequence alignment or GPCRdb.

It is worth noting that engagement of the agonists with the toggle switch generally induces structural rearrangement of the triad P^5.50^I^3.40^F^6.44^ motif in solved active GPCR structures^28,29^. Specifically, the shift of W^6.48^ caused a one-step downward shift of F^6.44^ in β2AR and A2AR, which allowed phenylalanine (F^6.44^) to fit into a hydrophobic pocket formed by I^3.40^ as the sidewall and the proline kink P^5.50^ at the bottom (Fig. 3d, Extended Data Fig. 7f). However, in GPBAR, the proline kink (P176^5.50^) moves away from F233^6.44^, which turns to interact with H107^3.47^. Instead, W237^6.48^ in GPBAR, which is in the position equivalent to F^6.44^ in other GPCRs, engages in hydrophobic interactions with L100^3.40^ and L103^3.43^ from TM3 and with V178^5.32^ from TM5 (Fig. 3d). Notably, the distance between W237^6.48^ and these leucines is larger than the distances between the traditional F^6.44^ vs. I^3.40^ pair in other receptors, and W237^6.48^-Y240^6.51^ creates a bulge at the helical turn in TM6 of GPBAR, which has not been described previously for any available GPCR structures (Fig. 3d). These structural features of GPBAR contribute to the loose contact between TM3 and TM6 and between TM5 and TM6.

Collectively, we conclude that the sensing of agonists by Y240^6.51^ and F96^3.36^, the shift of W237^6.48^ and the rearrangement of L100^3.40^ and V178^3.52^ might serve as the key molecular mechanisms of GPBAR activation, mimicking the role of “toggle-switch” and PIF motif, respectively, in the classical activation pathway of the typical class A GPCRs, and therefore connect the GPBAR ligand binding pocket to the G protein interaction site. These structural and functional studies imply that the toggle switch and the PIF motif derived from the sequence alignment may not always function according to the proposed mechanism of activation in a particular GPCR; the evolution of other key residues may substitute for the functions of these well-known residues through alternative structural combinations.

### Coupling to Gs through TM bundles and ICL3 of GPBAR

Due to the engagement of Gs with the extension of TM5 and TM6 and ICL3 between them, the α5 helix C-terminus of Gs does not penetrate as deeply into GPBAR as in other resolved Gs-coupled receptor complex structures (Fig. 4a, 4e). The recognition of the α5 helix of Gs by GPBAR involves TM3, TM5, TM6, ICL2 and ICL3. The resulting crevice is in general more hydrophilic compared with that in the β2AR**–**Gs complex and TM6 helix interacts with Gs more extensively (Extended Fig. 8a-c). Residues proximal to Gs have been confirmed by mutation experiments (Extended Fig. 8a-d and Table 4-5). The divergence of the G protein subtype at the Gs L394 and E392 positions may partly contribute to the selective coupling with Gs in preference to Gq by GPBAR (Extended Fig. 8e-f).

Outside the TM bundle, a unique feature of the GPBAR**–**Gs complex structure is the electron density covering the integral ICL3 (residues R201 to L214) that contributes to both Gs binding and activation (Extended Data Fig. 3c). The ICL3 of GPBAR forms three additional helical turns at the intracellular ends of TM5 and TM6 (in comparison with the active forms of β2AR or A2AR) and a bulge turn of approximately 6 residues between two helices. The more compact intracellular half of GPBAR brings these structures closer to Gs, leading to additional interaction at the C-terminal part of the Gα-Ras-like domain, including the β6, α4 and i3 loop (Fig. 4b)30. Importantly, three successive Arg, R201^ICL3^, R204^ICL3^ and R208^ICL3^, form charge interactions with the acidic patch produced by the i3 knob (Fig. 4b). These interactions, together with the hydrophobic packing of ICL3 of GPBAR with the α-helix 4 and the β-strand 6 of Gs, pull the i3 loop from T319 to D331 of Gs, corresponding to a shift of approximately 2 Å toward the receptor, inducing rearrangement of the α4-β6 turn and causing substantial side chain reorganization (Fig. 4b and Extended Fig.8g).

An inspection of the interactions between ICL3 (R201-L214) of GPBAR and Gs enabled us to deduce an R/KΨXR/KXΨXR motif that contributes to Gs recognition. Consistently, mutations of ICL3 residues of GPBAR, including R204A, V206A or R208A, significantly impaired P395**–**induced cAMP accumulation with respect to both potency and efficacy (Extended Fig.8h), thus confirming the importance of these specific residues in the ICL3 binding motif in Gs coupling (Fig. 4b). We then questioned whether the binding of the third intracellular loop of GPCRs to Gs is a common activation mechanism utilized by a subset of GPCRs, and therefore tried to screen out receptors sharing residue arrangement in the R/KΨXR/KXΨXR motif of ICL3 by sequence alignment (Fig. 4c). Sequence searching identified that at least 3 known Gs-coupled GPCRs, including V2R, PF2R and EP2 have a minimum of 2 corresponding residues in the R/KΨXR/KXΨXR motif. Moreover, we observed that mutations in corresponding motifs in the ICL3 regions of these receptors, significantly decreased Gs activation after engaging with their agonists (Fig. 4d).

### A putative second ligand binding pocket with allosteric properties

The high-quality cryo-EM density maps unveiled annular lipid molecules outside the seven transmembrane bundles in both INT-777– and P395–bound GPBAR signaling complexes (Extended Data Fig. 9a). These lipids are mostly found at the extracellular half of the receptor near the orthosteric binding pocket (Extended Data Fig. 9a). Most of these lipid binding sites are shallow indentations around the receptor surface, however, one unexpected but clear density in both cryo-EM density maps of GPBAR–Gs complexes were observed in the well-defined pocket constituted by TM3, TM4, TM5 and ICL2, where a similar lipid binding site for GPCR P2Y_1_ (PDB ID 4XNV)^31^ and an allosteric modulator site for GPR40^32^ have been reported (Fig. 5a and Extended Data Fig. 9b-c). We assigned a cholesterol into the electron density of second binding pocket of the P395–GPBAR–Gs complex (Fig.5a-b, and Extended Data Fig. 9c). For INT-777–GPBAR–Gs complex, both cholesterol and INT-777 could be fit into the same position. However, computational simulation indicated that both the GPBAR and the INT-777 bound at orthosteric site exhibit least RMSD fluctuations in the presence of the INT-777, but not the cholesterol, CHS or no ligand at this lipid binding site (Extended Data Fig. 9d-e). We therefore assigned the INT-777 at this lipid binding site and this assignment was further supported by following biochemical results (Fig.5e-g).

Notably, in both P395-bound– and INT-777-bound–GPBAR structures, the modeled cholesterol or INT-777 sits in a hydrophobic pocket and stabilizes the ICL2 in a loop-like conformation. Binding of a ligand at this site may release E109^3.49^ of the conserved D/ERY motif to recognize Y391 of Gs (Fig. 5b, Extended table 6). Importantly, mutations of the amino acids involving in the second binding sites, such as L104^3.44^ and L130^4.48^ to alanine, significantly impaired agonist -induced cAMP accumulation, whereas mutations of surrounding residues, such as the two Pro residues (P120G and P121G), had no significant effects (Fig. 5d, Extended table 4). These results suggest that a ligand bound to the second binding site might positively modulate the activation of GPBAR, which is likely due to further stabilizing the ICL2 in a conformation more readily for Gs coupling (Fig. 5b-5c, Extended table 6-8).

Considering INT-777 is a bile acid derivative, we suspected that INT-777 and other bile acids may be able to bind to this second pocket in GPBAR and that the bound bile acids may allosterically regulate receptor activity. We next screened all nine commercially available bile acids for their allosteric cooperativities. Notably, five of them, including CA, DCA, GCA, TCA and TDCA, showed modest but robust positive cooperative effects for GPBAR activation in response to the P399 interaction (Fig. 5e-5f and Extended table 9). Intriguingly, all five bile acids bearing allosteric properties contain a hydroxyl group substitution at position 12, whereas the other 4 bile acids do not, indicating a strong structural-function relationship (Fig. 5e). We next mutated all 8 residues surrounding the second bile acid binding site and test the positive cooperative effects using the five bile acids showing allosteric properties. Connecting to the orthosteric site, only the upper 4 residue mutations impaired the allosteric properties of all 5 bile acids (Fig. 5e and 5g). Importantly, the T131A mutation, which disrupted a potential H-bond between the 12-hydroxyl group of modeled INT-777 and other bile acid, abolished this positive cooperativity for all 5 bile acids (Fig. 5e,5g and Extended table 10). This observation is consistent with the observation that only bile acids bearing the 12-OH group exhibited allosteric functions. Taken together, these results demonstrated that the binding of bile acids bearing a 12-OH group to the second bile acid binding pocket of GPBAR has a positive allosteric effect on its orthosteric agonist binding and activity (Fig. 5d-5e).

### Structural basis of the biased property of INT-777

The arrestin-mediated GPBAR functions, which may contribute to the diverse signaling and cellular outputs elicited by different GPBAR ligands, have only recently begun to be appreciated^14^. Interestingly, the synthetic GPBAR agonist P395 was biased more heavily toward β-arrestin, with a β value of −0.46, whereas INT-777 displayed a bias property toward Gs, with a β value of 0.57, considering the endogenous bile acid LCA as a reference (Fig. 6a, Extended Data Fig. 10a). The β value was calculated through the operational model, which reflects the differences of both the efficacy and potency of two different pathways ^33,34^. Thus, a comparison of the INT-777–GPBAR complex structure with the P395–GPBAR complex structure could shed light on the structural basis of GPBAR signaling bias ^35^.

Although the overall structure of the INT-777–GPBAR complex is similar to that of the P395–GPBAR complex, the structural differences in specific residues may contribute to the signaling bias. Analysis of the root-mean-square deviation (RMSD) over Cα atoms between INT-777– and P395–bound GPBAR structures indicated that the most significant differences between INT-777– and P395–bound GPBAR structures were within the three extracellular loops and ICL1 (Fig. 6b, Extended Data Fig. 10b). We therefore performed alanine scanning mutagenesis of the residues with significant conformational differences (RMSD of Cα are larger than 2 Å) between the two structures and then examined the effects of the mutants on both Gs and arrestin downstream signaling. Mutation of Q77^ECL1^, P151^ECL2^, and P256^ECL3^ to alanine resulted in a significant decrease in arrestin recruitment that exceeded the decrease in cAMP accumulation (Fig. 6c-6d and Extended Data Fig. 10b-c). In addition to observations that were consistent with the previous finding that ECL3 in the GLP-1 receptor contributed to the bias property^36^, we found that ECL1 and ECL2 regions of GPBAR also contributed to arrestin-biased activation. In particular, the flipping of large side chains by Q77^ECL1^ and P151^ECL2^ and the correlated mutating effects on bias property changes indicated a potential structural-function relationship at these two extracellular loops. Another important conformational difference between INT-777– and P395–bound GPBAR was observed in ICL1 (Fig. 6b). Mutations of R44^ICL1^ and L45^ICL1^ to alanine significantly decreased arrestin recruitment but had little effect on Gs coupling (Fig. 6c-6d and Extended Data 10 b). Consistent with this finding, a direct interaction between ICL1 of GPBAR and Gs was not found in either structure.

Importantly, previous studies only identified that biased function of the exendin-P5–GLP-1R–Gs complex structure is mainly conferred by its increased Gs coupling activity without significant effects on arrestin coupling^36^. Conversely, our present results demonstrated that GPBAR gained biased properties through the regulation of arrestin activity without affecting Gs signaling, as mutations of Q77^ECL1^, P256^ECL3^ and R44^ICL1^L45^ICL1^ to Ala diminished arrestin recruitment without significantly affecting Gs activation (Extended Data Fig. 10b). The identified ECL1 and ECL3 regions important for biased signaling of GPBAR are more diverse than the previously identified ECL3 region for GLP-1R^36^. Furthermore, we anticipate that R44^ICL1^L45^ICL1^ in GPBAR could be the direct binding site of arrestin but not Gs. Therefore, our study supports the idea that biased signaling could be regulated through allosteric coupling of diverse regions from extracellular to intracellular portions.

## Discussion

The cryo-EM structures obtained in this study revealed a large oval pocket to accommodate the large steroid core of bile acids, sporadic hydrophilic residues on one side, along with hydrophobic residues on the opposite side, underlying the molecular mechanism of recognition of an ampholytic ligand by GPBAR. Moreover, key residues inside the orthosteric pocket are identified as important fingerprint readers to discriminate different bile acids with substitutions at the 7 (R1) and 12 (R2) positions and the conjugating groups at the C-termini of the steroid core. These specific interactions, as well as the identification of only bile acids with a structural feature of 12-OH substitutions to afford allosteric cooperative effects, may account for the different potencies and efficacies of bile acids in cAMP accumulation and diverse downstream functions through GPBAR activation.

Along and below the ligand binding pocket, there was an unusual separation of TM6 from central TM3, likely due to the absence of P^5.50^I^3.40^F^6.44^ motif packing in the GPBAR structure. This conserved packing functions to tether the TM3-TM5-TM6 bundles in other active GPCR structures (Fig. 4e). Notably, the structural rearrangement of the P^5.50^I^3.40^F^6.44^ and N^7.45^P^7.46^XXY^7.49^ motifs, as well as the shift of the “toggle switch” W^6,48^, are hallmarks for all known active class A GPCR structures determined to date ^28,29^. However, GPBAR does not contain the conserved NPXXY motif, and its TM bundles in the active state are linked by V178^5.52^L100^3.40^W237^6.48^ packing rather than tethering by the traditional P^5.50^I^3.40^F^6.44^motif, suggesting that diverse structural motifs exist among GPCRs to connect the ligand binding pocket to the G protein coupling site, despite their evolutionary closeness and similar key residues according to their Ballesteros-Weinstein numbers. Our mutagenesis and structural observations also suggested that the interactions among Y240, S247 and L166 form a potential hub for maintaining the inactive state of GPBAR, whereas engaging with Y240 with an H-bond and hydrophobic interactions provided by a ligand may induce both the Y240^6.51^ and W237^6.48^ switches to activate GPBAR.

In particular, both GPBAR**–**Gs complex structures revealed the coupling of GPBAR to the Gs protein through ICL3 of the receptor (Fig. 4e). In general, the function of the ICL3 region in GPCRs has not been defined, and no receptor**–**Gs complex structure has shown the integral electron density of the ICL3 to disclose its functions in effector coupling. Our structural analysis and biochemical study suggest that an R/KΨXR/KXΨXR motif in ICL3 could be a general mechanism utilized by a group of GPCRs to couple to Gs. These observations suggested that the coupling of ICL3 of GPCRs to G proteins could be important for effector activation in many cases, representing a mechanism that has not been previously recognized.

## ACKNOWLEDGEMENTS

We acknowledge support from the National Key Basic Research Program of China Grant 2018YFC1003600 to X.Y. and J.-P.S., the National Science Fund for Distinguished Young Scholars Grant (81773704 to J.-P.S. 81425024 to X.X), the National Science Fund for Excellent Young Scholars Grant (81822008 to X.Y., 81922071 to Y.Z.), Zhejiang Province National Science Fund for Excellent Young Scholars LR19H310001 to Y.Z., the National Natural Science Foundation of China Grant (31900936 to F.Y., 81730099 to X.X.) the China Postdoctoral Science Foundation Grant 2019T120587 to F.Y. and Innovative Research Team in University Grant IRT_17R68 (to X.Y. and Y.S.). Sample preparation for cryo-EM studies was supported by Protein Facility, Zhejiang University School of Medicine. The cryo-EM data were collected at the Center of Cryo-Electron Microscopy, Zhejiang University, with assistance of S. Chang and X. Zhang.

## AUTHOR CONTRIBUTIONS

Y.Z., X.X., X.Y., and J.-P.S. organized the whole project. J.-P.S., Y.Z., X.Y., X.X., supervised the overall project design and execution. Y.Z. guided all the Cryo-EM study. Y.Z. and J.-P.S guided all structural analysis. X.X. provided all ligand study and chemical guidance. X.Y. initiated the study of recognition mechanism of bile acid, allosteric assays and bile acid derivatives by GPBAR and designed the screening assay for complex formation. J.-P.S., X.Y.. designed all cellular experimental details. F.Y., L.L.G. and P.X. developed the GPBAR constructs and optimized protein expression. F.Y., L.L.G. and P.X. established P395/INT-777–GPBAR–Gs complex formation strategy; F.Y., L.L.G., P.X. and X.W. screened the bile acids or its derivatives for complex formation. F.Y., L.L.G., P.X. and X.W. performed virus production, insect cell expression and prepared samples for cryo-EM. F.Y., L.G. developed the method for solubilization of the bile acids advised by Y.Z., J.-P.S. and X.Y. F.Y., L.L.G. screened conditions for gel filtration. D.-D.S. evaluated the sample by negative-stain EM; C.Y.M. prepared the cryo-EM grids; C.Y.M., D.-D.S. collected the cryo-EM data with assistance from Q.Y.S.; C.Y.M. and Q.Q.M. performed cryo-EM map calculation, model building and structure refinement. C.M. assisted in protein purification in Protein Facility. L.L.G., P.X. and K.Z. performed pull down assay. X.Y., Y.Z., J.-P.S. and X.X. designed all the mutants for ligand binding pocket and second bile acid binding site. X.Y. designed the biased signaling assay. X.Y. and J.P.S. designed the experiments for characterization of bile acid binding patterns of GPBAR. X.Y. designed the cooperative assay for allosteric mechanism. Q.Y.S. carried out the computational simulations. J.Y.L. performed cooperative assay and data analysis. X.W., L.L.G., J.Y.L., S.M.G., L.Q.Z., F.Yi., Y.Q.P., X.Y.L. and K.Z. performed cAMP accumulation assay and binding assay. X.W., Y.Q.P., R.R.L. and S.M.G. performed BRET assay. Y.M.S., F.Yi., J.Y.Z. and C.T.J. participated in the design and explanation of the cAMP and BRET results and provided insightful ideas and experimental designs. V.C.L. oversaw the structural analysis. J.-P.S. wrote the manuscript. All the authors have seen and commented on the manuscript.

## METHODS

### Cryo-EM data acquisition

The purified P395–GPBAR–Gs complex (3.0 μL) at 4.0 mg/ml and INT-777–GPBAR–Gs complex (3.0 μL) at 4.5 mg/ml were applied onto a glow-discharged holey carbon grid (Quantifoil R1.2/1.3), and subsequently vitrified using a Vitrobot Mark IV (Thermo Fischer Scientific). Cryo-EM imaging was performed on a Titan Krios equipped with a Gatan K2 Summit direct electron detector in the Center of Cryo-Electron Microscopy, Zhejiang University (Hangzhou, China). The microscope was operated at 300kV accelerating voltage, at a nominal magnification of 29,000× in counting mode, corresponding to a pixel size of 1.014 Å. In total, 4826 movies of P395–GPBAR–Gs complex and 6229 movies of INT-777– GPBAR–Gs complex (1^st^, 4153 movies; 2^nd^, 2076 movies) were obtained at a dose rate of about 7.8 electrons per Å^2^ per second with a defocus range of −0.5 to −2.5 μm. The total exposure time was 8 s and intermediate frames were recorded in 0.2 s intervals, resulting in an accumulated dose of 62 electrons per Å^2^ and a total of 40 frames per micrograph.

### Image processing and 3D reconstruction

Dose-fractionated image stacks were subjected to beam-induced motion correction using MotionCor2.1^37^. A sum of all frames, filtered according to the exposure dose, in each image stack was used for further processing. Contrast transfer function (CTF) parameters for each non-dose weighted micrograph were determined by Gctf^38^. Particle selection, 2D and 3D classifications were performed on a binned dataset with a pixel size of 2.028 Å using RELION-3.0-beta2^39^. Semi-automated selected particles were subjected to reference-free 2D classification, producing particles with well-defined averages for further processing. The map of PTH1R**–**Gs complex (EMDB-0410) low-pass filtered to 20 Å was used as an initial reference model for 3D classification. Conformationally homogeneous subsets showed detailed features for all subunits were subjected to further 3D classification focusing the alignment on the complex with the exception of AHD of the Gαs, produced one stable subsets accounting for 185,911 and 92,816 particles for two datasets, respectively. The two datasets were subsequently combined and subjected to 3D refinement and Bayesian polishing and frames 1-20 were used in the final refinement to reduce background noise and improve EM map quality. The final map has an indicated global resolution of 3.0 Å at a Fourier shell correlation of 0.143. Local resolution was determined using the Bsoft package with half maps as input maps^40^.

### Model building and refinement

The initial homology model of GPBAR was generated using Phyre2. The β2 AR**–**Gs protein complex (PDB ID 3SN6) was to generate the initial models of Gs and Nb35.. Agonist and lipid coordinates and geometry restraints were generated using phenix.elbow^41^. Models were docked into the EM density map using UCSF Chimera^42^. This starting model was then subjected to iterative rounds of manual adjustment and automated refinement in Coot^43^ and Phenix^41^, respectively. The final refinement statistics were validated using the module ‘comprehensive validation (cryo-EM)’ in PHENIX^44^. Structural figures were prepared in Chimera, Chimera X^45^ and PyMOL (https://pymol.org/2/^46^). The final refinement statistics are provided in Extended Data Table 1.

